# Phenotypic variation in mitochondrial function across New Zealand snail populations

**DOI:** 10.1101/230979

**Authors:** Emma S. Greimann, Samuel F. Ward, James D. Woodell, Samantha Hennessey, Michael R. Kline, Jorge A. Moreno, Madeline Peters, Jennifer L. Cruise, Kristi L. Montooth, Maurine Neiman, Joel Sharbrough

**Author notes:** **CORRESPONDENCE TO:** Joel Sharbrough, Colorado State University, 440 Biology Building, Fort Collins, CO 80523.

## Abstract

Mitochondrial function is critical for energy homeostasis and should shape how genetic variation in metabolism is transmitted through levels of biological organization to generate stability in organismal performance. Mitochondrial function is encoded by genes in two distinct and separately inherited genomes – the mitochondrial genome and the nuclear genome – and selection is expected to maintain functional mito-nuclear interactions. Nevertheless, high levels of polymorphism in genes involved in these mito-nuclear interactions and variation for mitochondrial function are nevertheless frequently observed, demanding an explanation for how and why variability in such a fundamental trait is maintained. *Potamopyrgus antipodarum* is a New Zealand freshwater snail with coexisting sexual and asexual individuals and, accordingly, contrasting systems of separate vs. co-inheritance of nuclear and mitochondrial genomes. As such, this snail provides a powerful means to dissect the evolutionary and functional consequences of mito-nuclear variation. The lakes inhabited by *P. antipodarum* span wide environmental gradients, with substantial across-lake genetic structure and mito-nuclear discordance. This situation allows us to use comparisons across reproductive modes and lakes to partition variation in cellular respiration across genetic and environmental axes. Here, we integrated cellular, physiological, and behavioral approaches to quantify variation in mitochondrial function across a diverse set of wild *P. antipodarum* lineages. We found extensive across-lake variation in organismal oxygen consumption, mitochondrial membrane potential, and behavioral response to heat stress, but few global effects of reproductive mode or sex. Taken together, our data set the stage for applying this important model system for sexual reproduction and polyploidy to dissecting the complex relationships between mito-nuclear variation, performance, plasticity, and fitness in natural populations.

## INTRODUCTION

The production of ATP *via* cellular respiration is a critical component of eukaryotic function and fitness^1–5^, and the components of the oxidative phosphorylation (OXPHOS) pathway generally appear to be evolving under purifying selection^6–11^. Nevertheless OXPHOS genes are often polymorphic within species^3,12–15^, with important implications for phenomena ranging from mito-nuclear incompatibilities^16^ to mitochondrial introgression^17^ to DNA barcoding^18^. How mitochondrial haplotype is connected to phenotype and how mitochondrial function is maintained over evolutionary time despite mutational pressure^19^ remain unclear, despite extensive variation for metabolic and mitochondrial function in natural populations of a diverse array of species^3,20–27^. Although some variation in metabolic and mitochondrial traits has been linked to specific environmental correlates (*e.g.,* altitude^28,29^, temperature^30^, energy source^31^), we lack a systematic understanding of the genotype-phenotype-fitness map for traits that depend on mitochondrial function and how this map may vary across environmental gradients.

Teasing apart the sources of variation in mitochondrial function is further complicated by the genetic architecture underlying ATP production. The enzymes that carry out OXPHOS (excepting succinate dehydrogenase in animals, fungi, and most plants^32,33^) are encoded by genes that are partitioned across two separate cellular compartments (i.e., the nucleus and mitochondria) with different inheritance regimes (i.e., biparental vs. uniparental, respectively)^34^. The biology of these coevolving “mito-nuclear” interactions has been the subject of intense study^16,35–41^, but the consequences of coevolution between genomes for function and fitness have only been evaluated in a handful of species and for a handful of mito-nuclear genotypes^16,42–45^. This research has nevertheless yielded critical insights into how mito-nuclear genotypic variation produces phenotypic variation, and demonstrating that mito-nuclear epistasis can be a powerful force for shaping genetic variation in nature^46^. The separate inheritance of nuclear and mitochondrial variation during sexual reproduction presents a challenge for natural selection in shaping co-adapted mito-nuclear genotypes^47^. In particular, because nuclear genomes are biparentally inherited and mitochondrial genomes are uniparentally inherited, mitochondrial genomes will thus be swapped across diverse nuclear genomic backgrounds. This separate inheritance of nuclear and mitochondrial genomes can result in the loss of particular high-fitness mito-nuclear genotypes from populations, as selection favors the fixation of mitochondrial genomes that have the highest fitness on average across the suite of nuclear variants in the population^48^. Sex-linkage and sex-specific fitness effects comprise a particular set of conditions that may enable the maintenance of mito-nuclear variation in populations^49^. The overwhelming preponderance of sexual reproduction in nature would thus seem to imply that the reliable transmission of particular mito-nuclear combinations is not all that important. What is perhaps more likely is that we do not have a good understanding of how sexual reproduction affects the connection between mito-nuclear genotype and mitochondrial performance.

Asexual reproduction, in which mitochondrial and nuclear genomes are co-inherited from mother to daughter, offers a useful contrast for understanding the consequences of sexual reproduction for mitochondrial function. In particular, mitochondrial genomes that are “trapped” in lineages that have transitioned to asexuality are inextricably linked to their nuclear genomic background for the remainder of their existence. Because asexual lineages are expected to accumulate deleterious mutations in both the nuclear and mitochondrial genomes^50^ as a result of Muller’s Ratchet^51^ and the Hill-Robertson Effect^52^, asexual lineages should exhibit reduced mitochondrial performance relative to sexual lineages. However, co-inheritance of nuclear and mitochondrial genomes also allows epistatic variation to become visible to natural selection, such that asexual lineages may experience both relatively intense and efficient selection favoring compensatory co-evolution between nuclear and mitochondrial loci. The extent to which asexual lineages exhibit reduced vs. improved mitochondrial performance compared to close sexual relatives will provide a useful guide for whether and how epistatic mito-nuclear variation contributes to mitochondrial performance.

*Potamopyrgus antipodarum* is a New Zealand freshwater snail^53^ featuring frequent co-existence and competition among obligately sexual and obligately asexual lineages^54^. So similar are these sexual and asexual lineages that the only reliably distinguishable character apart from reproductive mode is that asexual lineages are polyploid and sexual lineages are diploid^55^. Because asexuality has arisen on multiple separate occasions within *P. antipodarum*^56,57^, distinct asexual lineages can be treated as repeated “natural experiments” into the consequences of sexual reproduction for mitochondrial performance^8,58^. *Potamopyrgus antipodarum* populations also feature extensive population structure in mitochondrial DNA sequences^8,56,59^, and asexual lineages harbor heritable variation for mitochondrial performance^27^. Notably, the occasional production of male offspring by obligately asexual female *P. antipodarum*^60^ makes it possible to evaluate the phenotypic consequences of mitochondrial mutations that are neutral or beneficial when present in females but deleterious when present in males. In particular, selection against male-harming mutations is expected to be relaxed in asexual lineages, whose mitochondrial genomes have been “trapped” in females for generations, such that asexual males are predicted to exhibit especially poor mitochondrial performance. Together, these features make *P. antipodarum* uniquely well suited for decoupling sexual reproduction from mitochondrial genome inheritance, a necessary step if we hope to understand how mito-nuclear epistasis contributes to organismal function and fitness in natural populations.

The observation that mitochondrial genomes of asexual *P. antipodarum* lineages accumulate putatively harmful mutations more readily than do their sexual counterparts^8,58^, combined with the heritable variation for mitochondrial function present in asexual lineages^27^ leads to the expectation that asexual *P. antipodarum* will exhibit reduced mitochondrial performance compared to sexual lineages. Here we tested that prediction as well as whether sex (*i.e.*, male vs. female) and source population affect mitochondrial and physiological function at organellar and organismal levels under laboratory conditions using wild-caught *P. antipodarum*.

## MATERIALS AND METHODS

### Field collections of *P. antipodarum*

The phenotypic and ecological similarity of sexual *vs.* asexual *P. antipodarum* enables direct comparisons across reproductive modes, but also means that definitive determination of reproductive mode (via flow cytometric genome size estimation – see below) requires snail sacrifice. We therefore sampled field-collected snails from New Zealand lakes known to harbor both sexual and asexual individuals^57,59,61^. Snails were collected from New Zealand lakes in January 2015 and December 2016 and transported by hand in damp paper towels to the University of Iowa. Upon arrival, snails were housed at 16°C with a 18L:6D photoperiod and fed *Spirulina* algae three times per week, as previously described^62^. We arbitrarily selected adult snails from lake collections (each of which consisted of hundreds to thousands of individuals) and isolated each snail in a 0.5 L glass container with 300ml carbon-filtered H_2_O. Water was changed weekly. All functional assays began immediately following isolation and were completed within six months of arrival at the University of Iowa.

To maximize our ability to sample and compare sexual and asexual individuals from the same lake, we assayed oxygen consumption under heat stress (details below) in 158 wild-caught females from six lakes (Figure 1, Table 1) and used destructive sampling to determine reproductive mode in snails that survived all three temperature trials. The recent discovery of asexually produced males in *P. antipodarum*^60^ and recognition of their unique potential to provide insight into the capacity for mtDNA to harbor mutations that are deleterious in males but neutral or beneficial in females motivated us to include both males and females in our next experiment. Here, we assayed behavioral function in response to heat stress and mitochondrial membrane potential in 46 wild-caught snails (Figure 1, Table 1).

**Table 1.** Summary of source populations of *Potamopyrgus antipodarum* sampled from New Zealand lake collections.

| **Oxygen consumption assay** |  |  |  |  |  |  |
| --- | --- | --- | --- | --- | --- | --- |
| **Lake** | **Latitude, Longitude** | **Sexual** | **Asexual** | **N/A<sup>1</sup>** | **Male** | **Female** |
| Alexandrina | -43.900476, 170.453978 | 12 | 3 | 11 | - | 26 |
| Clearwater | -43.602131, 171.043917 | - | 3 | 17 | - | 20 |
| Kaniere | -42.832886, 171.14759 | 16 | - | 12 | - | 28 |
| Paringa | -43.713068, 169.411348 | 4 | - | 20 | - | 24 |
| Rotoroa | -41.855414, 172.637882 | - | 16 | 22 | - | 38 |
| Selfe | -43.237765, 171.520449 | - | 3 | 19 | - | 22 |
| **Total** | - | **32** | **25** | **101** | **0** | **158** |
| **Behavior and mitochondrial membrane potential assays<sup>2</sup>** |  |  |  |  |  |  |
| **Lake** | **Latitude, Longitude** | **Sexual** | **Asexual** | **N/A** | **Male** | **Female** |
| Alexandrina | -43.900476, 170.453978 | 3 | - | - | 3 | - |
| Ellery | -44.046898, 168.654261 | 2 | 3 | - | - | 5 |
| Kaniere | -42.832886, 171.14759 | 5 | 1 | - | 4 | 2 |
| Mapourika | -43.315212, 170.204061 | 8 | 2 | - | 6 | 4 |
| Rotoroa | -41.855414, 172.637882 | 4 | 1 | - | - | 5 |
| Selfe | -43.237765, 171.520449 | 9 | 8 | - | 9 | 8 |
| **Total** | - | **31** | **15** | **0** | **22** | **24** |

<sup>1</sup> – N/A refers to snails that did not survive all three temperature trials <sup>2</sup> – Same individual snails were used in behavioral and mitochondrial membrane potential assays

**Figure 1.**
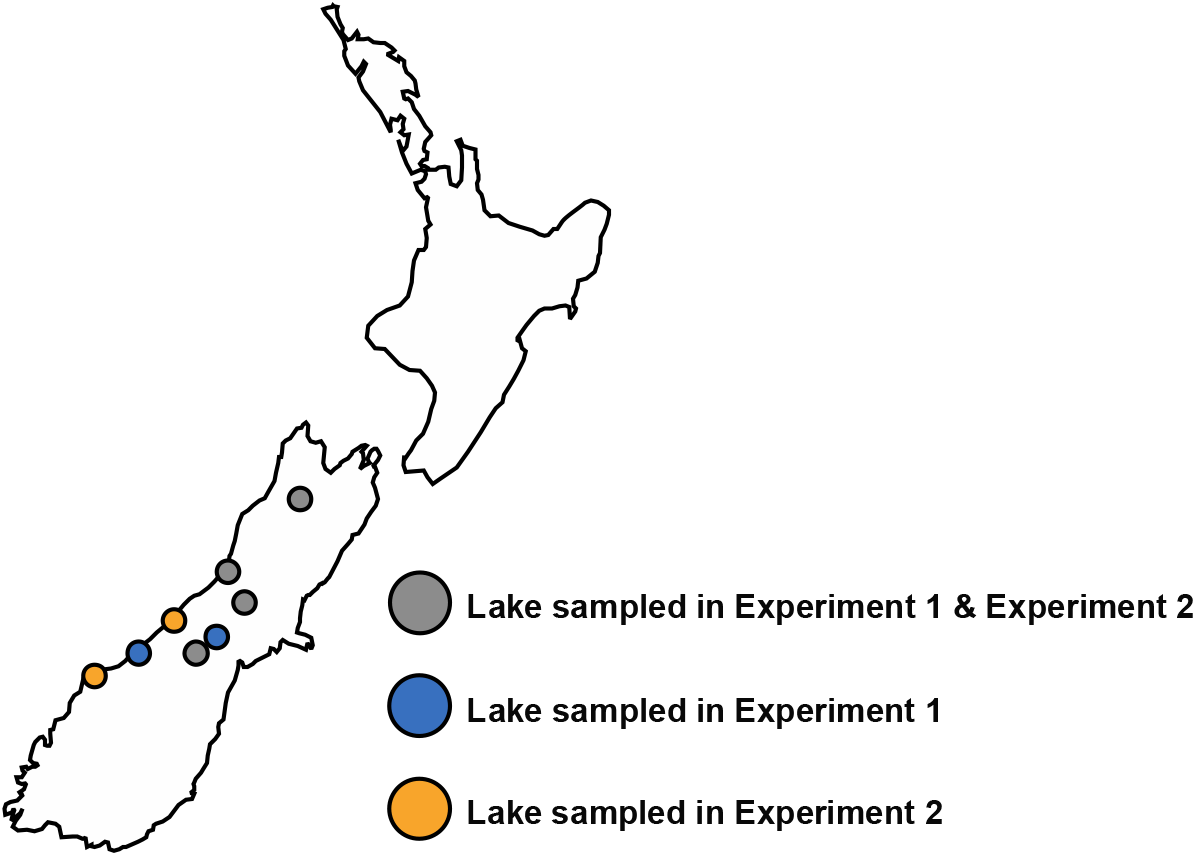
Sampling *P. antipodarum* from New Zealand lakes. Samples from four lakes were used in both experiments (grey circles – Lakes Alexandrina, Kaniere, Rotoroa, and Selfe), samples from two lakes were used in the oxygen consumption experiment only (blue circles – Lakes Clearwater and Paringa), and samples from a different two lakes were used for the behavioral and mitochondrial membrane potential experiments only (orange circles – Lakes Ellery and Mapourika).

### Oxygen consumption under heat stress conditions

Because oxygen becomes limiting to ectotherms under elevated temperatures^63^ and because *P. antipodarum* demonstrates signs of stress at elevated (~30°C) temperatures (*e.g.,* reduced fecundity^64^, elevated oxygen consumption^27^, and decreased righting ability^27^), we measured oxygen consumption using an aquatic respirometer as previously described^27^ for 158 wild-caught female *P. antipodarum* from each of six lakes. Oxygen consumption was assayed at three different water temperatures: 16°C (not stressful, and similar to New Zealand lake temperatures), 22°C (moderately stressful), and 30°C (stressful)^27^. Each snail was assayed at each temperature in a randomly determined order, and only the 61 snails (38.6%) that survived across all three temperature treatments were included in our analyses. We were unable to determine a single source of mortality for those snails that did not survive all three temperature trials, although exposure to high temperatures combined with hypoxic or near-hypoxic conditions inside the respiration chamber may certainly have played a role. *Potamopyrgus antipodarum* are also notoriously difficult to keep alive in laboratory conditions when isolated from other snails (personal observation), and our temperature treatment procedure combined with recovery periods required that snails spent many weeks in isolation. Wet mass for each individual was calculated after each trial, and mean mass across all three trials was used in our final analyses.

### Behavioral response to heat stress

Righting time and time to shell emergence following a startling stimulus increase with temperature in *P. antipodarum*^27^, indicating that both assays can be used to assess heat-induced stress. We quantified righting times and time to emergence under each of the same three temperature treatments used for oxygen consumption in 46 wild-caught *P. antipodarum* and compared behavior reaction norms across temperatures, lakes, reproductive modes, and sexes. Snails were each assayed once at each temperature, and no snails died in the course of behavioral analyses, which require ~50% less time spent (<1.05 hr) under the elevated temperature treatment than oxygen consumption assays (2 hr) and no time spent under hypoxic or near-hypoxic conditions.

### Mitochondrial membrane potential

JC-1 is a small positively charged molecule that diffuses down the electrochemical gradient across the inner mitochondrial membrane^65^. Under UV illumination, JC-1 fluoresces green if dispersed and red if aggregated (e.g., when inside the mitochondrial matrix)^65^. As a result, the median ratio of red: green fluorescence in freshly isolated live mitochondria stained with JC-1 can serve as a proxy for mitochondrial membrane potential, with a high ratio indicating high membrane potential. We isolated mitochondria from 46 wild-caught *P. antipodarum* and measured red: green ratios in JC-1-treated mitochondrial extracts as previously described^27^ using a Becton Dickenson LSR II flow cytometer. Using the set of 46 snails that completed behavioral assays at all three temperatures, we compared median red: green JC-1 ratios across lakes, reproductive modes, and sexes.

### Determination of reproductive mode

Sexual *P. antipodarum* are diploid and asexual *P. antipodarum* are polyploid (triploid or tetraploid^57,66^), allowing us to use flow cytometry to determine DNA content and infer reproductive mode of individual snails. After completing all physiological and/or behavioral assays blind with respect to snail reproductive mode, we dissected head tissue from each individual snail, flash froze this tissue in liquid nitrogen, and stored the frozen tissue at −80°C until flow cytometry. We then homogenized head tissue in DAPI solution, passed this homogenate through a 30 μm filter, and analyzed the suspended nuclei on a Becton-Dickinson FACS Aria II following the protocol (including a chicken red blood cell standard) outlined in previous work^67^.

### Statistical analyses

Using a mixed-effects model, we tested whether mass, temperature (16°C, 22°C, 30°C), lake-of-origin (n = 6), reproductive mode (sexual or asexual), and interactions of mass × temperature, mass × lake, mass × reproductive mode, temperature × lake-of-origin, temperature × reproductive mode, mass × temperature × lake, and mass × temperature × reproductive mode were good predictors of oxygen consumption per hour. A term for snail identity was fit as a random intercept to account for repeated measures on individuals across temperatures. We employed a similar framework for emergence time following a startling stimulus and righting behavior after being flipped upside-down, except we also included terms for sex (male or female) and a temperature × sex interaction, as we had both male and female snails in the sample used in these analyses. Finally, we modeled mitochondrial membrane potential, measured as the ratio of red: green fluorescence, as a function of lake, reproductive mode, and sex, using analysis of variance (ANOVA) and a Levene’s test to check for equal variances (*F*_5,40_ = 2.044, *p* = 0.09). Because we lacked replication of reproductive mode and sex within all sampled lakes, and after discovering a large lake effect in several phenotypic measures, we also fit sub-models predicting each phenotypic measure within lakes. Sub-models were equivalent to those described above for phenotypic measures but only included data from lakes that had replication for reproductive mode, sex, or both.

We developed final models using backwards selection until only predictors with *p*-values less than 0.05 remained. All main effects were kept in the model when part of a significant interaction. To test assumptions of normality and heteroscedasticity of errors, we graphically inspected residuals and log- or power-transformed response variables when necessary (Figure S3). Power transformation parameters were estimated using the geoR package^68^. We performed all statistical analyses in R^69^, fitting fixed-effect models with the *lm* function, fitting mixed-effects models using the lme4 package^70^, estimating degrees of freedom for mixed-effect models using Satterthwaite’s approximation *via* the lmerTest package^71^, and performing pairwise comparisons of mixed model predictors using the emmeans package^72^, which adjusts *p* values for multiple comparison using the Tukey method. To test whether behavioral phenotypes were correlated with mitochondrial membrane potential, we estimated pairwise Pearson correlation coefficients for behavioral responses at each temperature and membrane potentials using the Hmisc package^73^ and corrected for multiple comparisons using the Holm procedure^74^.

## RESULTS

### Prevalence of reproductive modes and sexes in two samples of wild-caught *P. antipodarum*

Our first sample was comprised of 158 female *P. antipodarum* collected from six New Zealand lakes (Lakes Alexandrina, Clearwater, Kaniere, Paringa, Rotoroa, and Selfe [Figure 1]), among which we were able to assay oxygen consumption at 16°C, 22°C, and 30°C for 61 snails. We determined ploidy (and therefore reproductive mode) for 57 out of these 61 snails. The sample included 32 sexual snails, 25 asexual snails, and four snails of unknown reproductive mode (Table 1). All 32 sexuals were diploid and all 25 asexual individuals were triploid. Only one of our sampled lakes included multiple sexual and asexual snails (Lake Alexandrina), while the other five were monomorphic for reproductive mode (Table 1).

Our second sample was comprised of 22 male and 24 female *P. antipodarum* collected from six New Zealand freshwater lakes (Lakes Alexandrina, Ellery, Kaniere, Mapourika, Rotoroa, and Selfe – Figure 1), among which we assayed emergence time and righting behavior at 16°C, 22°C, and 30°C, quantified mitochondrial membrane potential, and determined ploidy for all 46 snails. This sample included 31 sexuals and 15 asexuals, and we sampled multiple sexual and asexual individuals from within three different lakes (Lakes Ellery, Mapourika, and Selfe). Lakes Mapourika and Selfe also had replication for sex, with both male and female sexuals sampled from both lakes. However, we were only able to sample male and female snails exhibiting both sexual and asexual reproduction in Lake Selfe (Table 1).

### Oxygen consumption under stressful conditions

To partition variation in oxygen consumption across biological and environmental axes, we measured snail oxygen consumption in a closed-system aquatic respirometer under three different temperature treatments. Individual snail reaction norms from each of six lakes for oxygen consumption across temperature treatments are depicted in Figure S1a. We fit a mixed-effects model to our response variable of oxygen consumption per hour. After eliminating non-significant predictors using backwards selection, we found significant effects of snail wet mass (*χ*^2^ = 10.82, df = 1, *p* = 0.0010), temperature treatment (*χ*^2^ = 45.91, df = 2, *p* < 0.0001), and lake-of-origin (*χ*^2^ = 12.74, df = 5, *p* = 0.0260) (Table 2). Snail wet mass had a positive effect on oxygen consumption per hour (correlation slope estimate = 5,859.06), indicating that larger snails consume more oxygen per hour than do smaller snails (Figure S2). *Post hoc* pairwise t-tests revealed that snails consumed significantly more oxygen at 22°C than at 16°C (*t* ratio = − 4.12, df = 118, *p* = 0.0002), at 30°C than at 16°C (*t* ratio = −6.72, df = 120, *p* < 0.0001), and at 30°C than at 22°C (*t* ratio = −2.663, df = 118, *p* = 0.0238) (Figure 2a). Similarly, *post hoc* pairwise t-tests of reaction norms across lakes revealed two statistical groups of lakes, with Lake Clearwater exhibiting especially low rates of oxygen consumption per hour. We did not find any difference in oxygen consumption per hour across reproductive modes (Figure 3a), even when we restricted our analyses to snails within Lake Alexandrina (Table S1, model e), the one lake where we had the snails required to perform such comparisons. Overall, these data indicate that lake-of-origin plays a major role in how snails respire in response to heat stress, and that asexual snails do not exhibit altered oxygen consumption compared to sexual snails overall. Still, low levels of within-lake sampling precluded a rigorous test of the effects of reproductive mode on oxygen consumption in response to heat stress.

**Table 2.** Fixed-effect only and mixed-effects models of select predictors on oxygen consumption, righting time, emergence time, and mitochondrial membrane potential.

| Oxygen consumption<sup>1</sup> |  |  |  |  |  |  |
| --- | --- | --- | --- | --- | --- | --- |
| Model | Factor | $\chi^2$ | df | *p* | Non-significant predictors<sup>2</sup> | |
| a | Intercept<sup>3</sup> | 5.32 | 1 | 0.0211 | Lake-of-Origin × Mass × Temperature; Mass × Reproductive Mode × Temperature; Lake-of-Origin × Mass; Mass × Temperature; Mass × Reproductive Mode; Reproductive Mode × Temperature; Reproductive Mode; Lake-of-Origin × Temperature | |
|  | Mass | 10.82 | 1 | 0.0010 |  |  |
|  | Temperature | 45.91 | 2 | < 0.0001 |  |  |
|  | Lake-of-Origin | 12.74 | 5 | 0.0260 |  |  |
| Emergence Time<sup>4</sup> |  |  |  |  |  |  |
| Model | Factor | $\chi^2$ | df | *p* | Non-significant predictors | |
| b | Intercept<sup>3</sup> | 138.14 | 1 | < 0.0001 | Reproductive Mode × Sex × Temperature; Sex × Temperature; Lake-of-Origin × Temperature; Reproductive Mode × Temperature; Reproductive Mode × Sex; Reproductive Mode; Sex | |
|  | Temperature | 48.53 | 2 | < 0.0001 |  |  |
|  | Lake-of-Origin | 11.34 | 5 | 0.0450 |  |  |
| Righting time<sup>5</sup> |  |  |  |  |  |  |
| Model | Factor | $\chi^2$ | df | *p* | Non-significant predictors | |
| c | Intercept<sup>3</sup> | 238.80 | 1 | < 0.0001 | Reproductive Mode × Sex × Temperature; Reproductive Mode × Temperature; Sex × Temperature; Lake-of-Origin × Temperature; Lake-of-Origin | |
|  | Temperature | 73.36 | 2 | < 0.0001 |  |  |
|  | Reproductive Mode | 4.48 | 1 | 0.0343 |  |  |
|  | Sex | 2.49 | 1 | 0.12 |  |  |
| | Reproductive Mode × Sex | 6.52 | 1 | 0.0107 | | |
| Mitochondrial membrane potential<sup>6</sup> |  |  |  |  |  |  |
| Model | Factor | Sum of Squares | df | *F* | *p* | Non-significant predictors |
| d | Intercept | 3.911 | 1 | 64.53 | < 0.0001 | Reproductive Mode × Sex; Reproductive Mode; Lake-of-origin |
|  | Sex | 0.42 | 1 | 6.84 | 0.0122 |  |
|  | Residuals | 2.67 | 44 |  |  |  |

<sup>1</sup> – Type III Repeated-Measures Analysis of Deviance $\chi^2$ Test of O<sub>2</sub> consumption per hour per gram
<sup>2</sup> – Semi-colon delimited list of non-significant predictors in order of elimination from the model (i.e., worst predictor listed first)
<sup>3</sup> – Snail ID was fit as a random intercept
<sup>4</sup> – Type III Repeated-Measures Analysis of Deviance $\chi^2$ Test of Box-Cox-transformed emergence time ( $\lambda = -0.2845$ )
<sup>5</sup> – Type III Repeated-Measures Analysis of Deviance $\chi^2$ Test of log-transformed righting times
<sup>6</sup> – Type III Analysis of Variance *F* Test of log-transformed ratios of red: green in mitochondrial extracts

**Figure 2.**
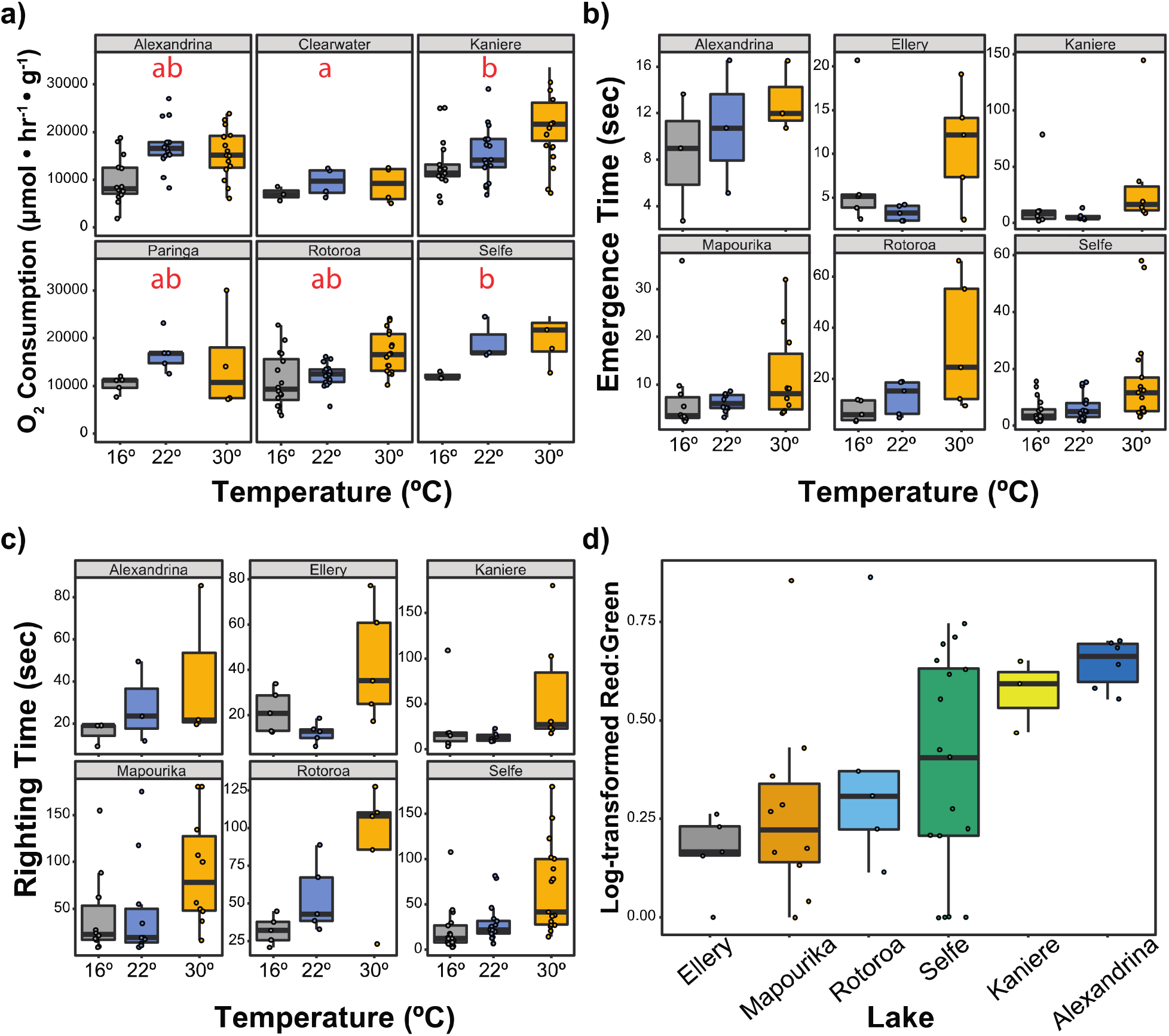
Mitochondrial phenotypic variation across New Zealand lake populations of *P. antipodarum*. **a)** Oxygen consumption/hour/gram measured at 16°C (gray), 22°C (blue), and 30°C (orange) in snails from six New Zealand lakes. There was a significant effect of mass, temperature, and lake-of-origin interaction on O_2_ consumption (Table 2a). Red letters indicate *post hoc* statistical groupings. **b)** Time to emergence following a startling stimulus was measured at 16°C (gray), 22°C (blue), and 30°C (orange) in snails from six New Zealand lakes. There was a significant effect of the temperature and lake-of-origin on emergence time (Table 2b). **c)** Time required for righting after being flipped was measured at 16°C (gray), 22°C (blue), and 30°C (orange) in snails from six New Zealand lakes. There was a significant effect of temperature, reproductive mode, and reproductive mode × sex interaction on righting time (Table 2c). **d)** Log-transformed ratios of red: green fluorescence following treatment with JC1 were measured using flow cytometry. There was a significant effect of sex on red: green ratios (Table 2d).

**Figure 3.**
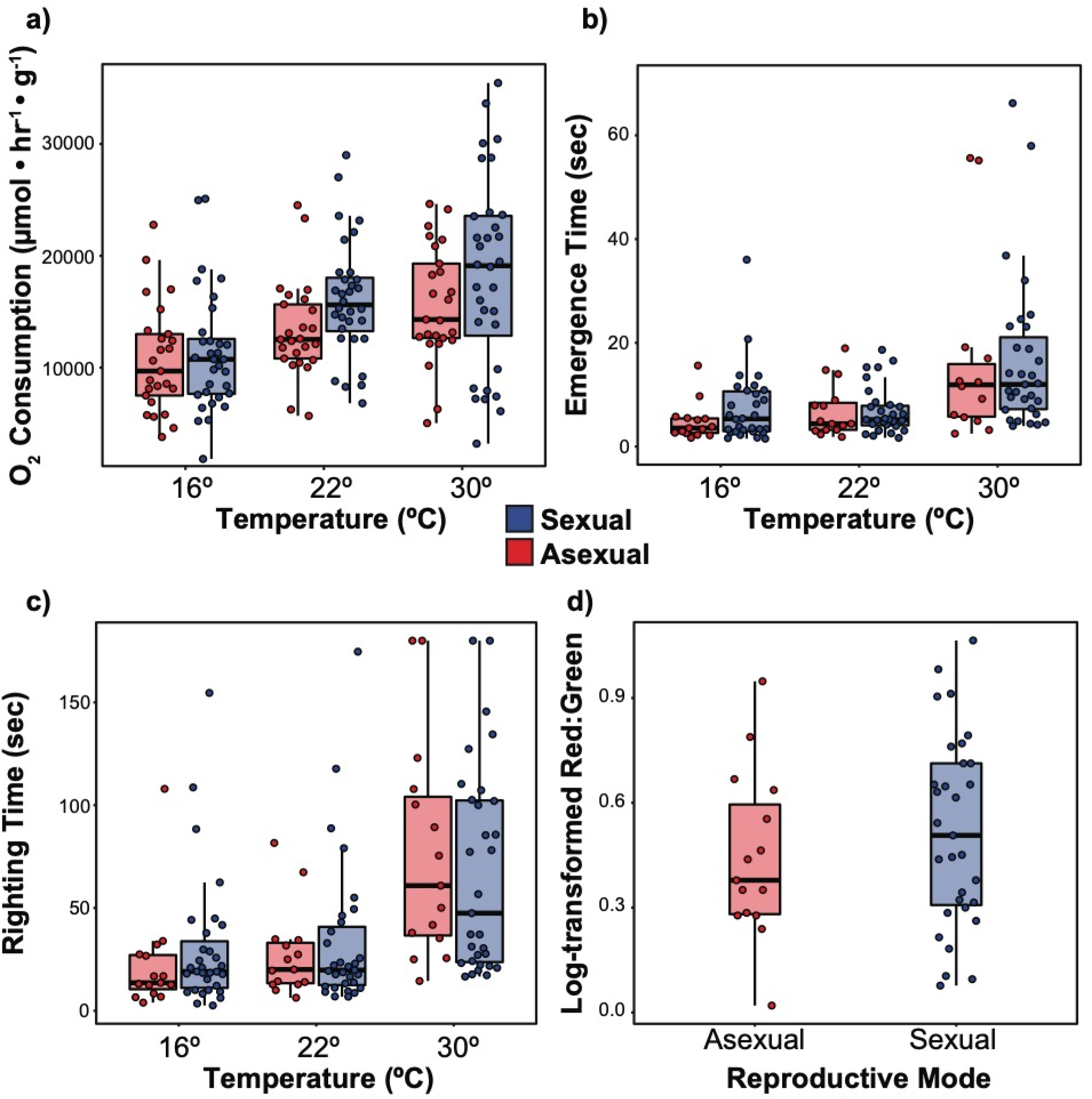
Mitochondrial phenotypic variation across reproductive mode among field-collected *P. antipodarum*. Measurements collected from sexual individuals are depicted in blue and measurements collected from asexual individuals are depicted in red. **a)** Oxygen consumption per hour per gram assayed at 16°C, 22°C, and 30°C. **b)** Time to emergence following a startling stimulus after incubation at 16°C, 22°C, and 30°C. Two outliers are not shown for the sake of display. **c)** Righting time following being incubated at 16°C, 22°C, and 30°C, and subsequently flipped over. **d)** Log-transformed ratios of red: green following treatment with JC1 are depicted in sexual vs. asexual individuals.

### Behavioral responses to heat stress

Using our second sample of 46 male and female *P. antipodarum* from six New Zealand lakes, we assayed two behaviors that challenge snails, especially under heat stress conditions: emergence time following a startling stimulus and righting ability after being flipped upside down. Individual snail reaction norms from each of six lakes for emergence time and righting time across temperature treatments are depicted in Figure S1b and c, respectively. We found that both temperature (*χ*^2^ = 48.53, df = 2, *p* < 0.0001) and lake-of-origin (*χ*^2^ = 11.34, df = 5, *p* = 0.0450) were significant predictors of emergence time (Table 2). In particular, *post hoc* t-tests of temperature effects averaged across lakes revealed that snails emerged from their shells significantly faster at 30°C than at either 16°C (*t* ratio = 6.42, df = 90, *p* < 0.0001) or 22°C (*t* ratio = 5.55, df = 90, *p* < 0.0001), but that there was no significant difference in emergence time at 16°C vs. 22°C (*t* ratio = 0.87, df = 90, *p* = 0.66) (Figure 2b). Despite the overall effect of lake-of-origin on emergence time, *post hoc* pairwise comparisons of lakes averaged over temperature did not reveal any obvious statistical groupings (*p* > 0.05 for all pairwise comparisons). We did not detect overall effects of reproductive mode (Figure 3b) or sex (Figure 4a) on emergence time, even when we limited our analyses to Lake Selfe (the only lake with replication for both sex and reproductive mode) (Table S1, model f). Temperature was also a significant predictor of righting behavior (*χ*^2^ = 73.36, df = 2, *p* < 0.0001), and *post hoc* pairwise analyses revealed that snails take significantly longer to right themselves 30°C than at 16°C (*t* ratio = 6.42, df = 90, *p* < 0.0001) and at 22°C (*t* ratio = 6.42, df = 90, *p* < 0.0001), but not at 22°C vs. 16°C (*t* ratio = 6.42, df = 90, *p* < 0.0001) (Table 2). In contrast to emergence time, reproductive mode (χ^2^ = 4.48, df = 1, *p* = 0.0343) and the reproductive mode × sex interaction (χ^2^ = 12.67, df = 2, *p* = 0.0107) were both significant predictors of righting time (Table 2), with asexuals righting themselves faster at 16°C and 22°C, but taking longer to right themselves at 30°C than sexuals (Figure 3c). Lake-of-origin was not a significant predictor of righting time overall (Figure 2c), but there were significant differences among lakes (*χ*^2^ = 12.67, df = 2, *p* = 0.0018) as well as a significant temperature × lake-of-origin interaction (*χ*^2^ = 12.99, df = 4, *p* = 0.0113) when we limited our analyses to females (Table S1, model s). We also tested whether behavioral responses were correlated with one another and found that significant negative correlations exist between emergence time and righting time at 16°C (*r*^2^ = 0.56, *p* < 0.0001), 22°C (*r*^2^ = 0.34, *p* < 0.0001), 30°C (*r*^2^ = 0.19, *p* < 0.0023), and between emergence time at 22°C and righting time at 16°C (*r*^2^ = 0.049, *p* < 0.0044). Together these data support a major role for lake-of-origin in determining behavioral response to heat stress, and we see some evidence from this relatively small set of snails that reproductive mode and sex have effects on righting behavior but not emergence.

**Figure 4.**
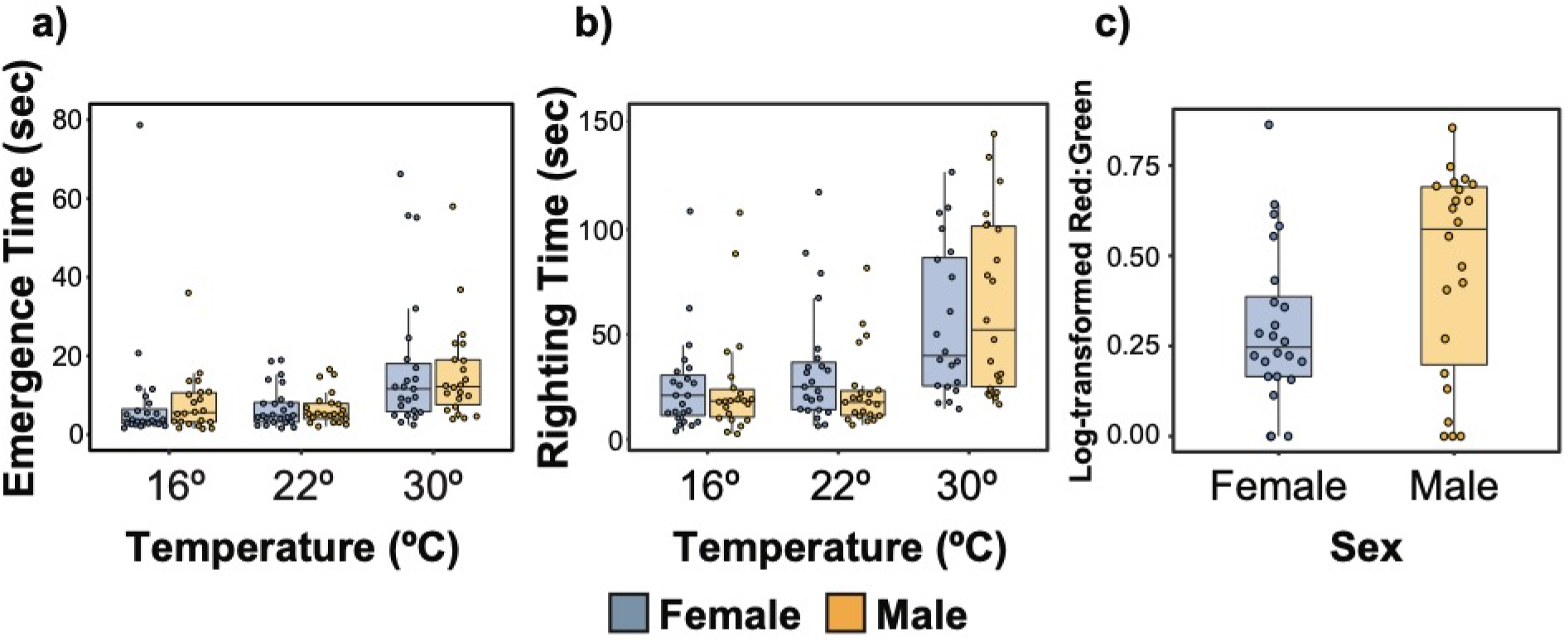
Mitochondrial phenotypic variation across sexes among field-collected *P. antipodarum*. Measurements collected from females are depicted in blue and measurements collected from males are depicted in yellow. **a)** Time to emergence following a startling stimulus. **b)** Righting time after being flipped, **c)** Log-transformed ratios of red: green following treatment with JC1.

### Mitochondrial membrane potential

We next assayed mitochondrial membrane potential in freshly isolated mitochondrial fractions, using the fluorescent dye JC-1, in this same set of 46 field-collected snails that were used for behavioral trials. We found that sex was a significant predictor of mitochondrial membrane potential (ANOVA *F*_1,44_ = 6.84, *p* = 0.0122, Figure 4c), with males having higher mitochondrial membrane potential than females. This difference between males and females was evident even when we limited our analysis to asexuals (ANOVA *F*_1,6_ = 12.09, *p* = 0.0132, Table S1, model k) or to Lake Selfe, which featured full replication of reproductive mode and sex (ANOVA *F*_1,15_ = 6.58, *p* = 0.0216, model h). While sexual males have higher mitochondrial membrane potentials than sexual females, the difference is not statistically significant (Table S1, model u), indicating that the higher membrane potential of asexual males are largely responsible for the effect of sex in the overall dataset. Lake-of-origin (Figure 2d) and reproductive mode did not affect mitochondrial membrane potential (Figure 3d), even when we limited our analyses to Lake Selfe or to females within Lakes Mapourika and Selfe (Table S1). We did not find evidence of correlations between mitochondrial membrane potential and either of the behavioral responses from the same snails. In sum, we found a strong effect of sex, especially in the context of asexuality, and no evidence of an effect of reproductive mode on mitochondrial membrane potential, with the caveat that low-within lake sampling may prevent us from observing relatively small effects.

## DISCUSSION

Extensive work in the *P. antipodarum* system has documented lake-specific phenotypes and local adaptation for resistance to infection by the trematode parasite *Atriophallophorus winterbourni* ^61,75–81^, life history traits such as growth rate and size^82^, and response to nutrient limitation^67,83^. Here, we report the first evidence of lake-structured variation for metabolic and behavioral function and sex-specific mitochondrial function in *P. antipodarum*. Combined with marked population structure for mitochondrial genetic variation^8,56^, this result suggests the intriguing possibility that mitochondrial function is locally tuned in *P. antipodarum*.

While the elevated linkage disequilibrium expected to result from asexuality should reduce the efficacy of natural selection in both nuclear^52,84–88^ and mitochondrial genomes^19,50,89^, stable transmission of mito-nuclear genotypes may also facilitate rapid mito-nuclear coadaptation and thereby local adaptation^47^. Surveys of mitochondrial genomes of asexual lineages^90–92^, including in *P. antipodarum*^8,56^, have revealed elevated rates of putatively harmful mutations in mitochondrial genomes compared to sexual lineages. Absent nuclear compensation for mitochondrial function^37,93^, we predict mitochondrial genomes carried by asexual lineages will therefore be associated with reduced mitochondrial performance. However, we could not detect any global decrease in mitochondrial, organismal, or behavioral performance in asexual vs. sexual lineages in this geographically diverse sample. We must therefore conclude that either the decline in mitochondrial performance in asexual lineages is only detectable among closely related sexual and asexual pairs within the same lakes or that asexual lineages are compensating for the increased mutation load in their mitochondrial genomes in some way. This compensation could be through mechanisms of physiological homeostasis (see e.g.^94^) or through clonal selection favoring lineages with mito-nuclear genotypes in which mutations in the nuclear genome compensate for the deleterious mutations in the mitochondrial genome. One potentially important source for genetic and/or physiological compensation might be polyploidy, as sexual *P. antipodarum* are all diploid and asexual *P. antipodarum* are all polyploid^55^. Extra genome copies in the nuclear genome may allow for mutational masking that facilitates broader traversal of the adaptive landscape^95^. Physiologically, elevated genome size is positively correlated with larger cell size^96^, which likely alters the signaling, energetic, and stoichiometric landscape of the cell^95^. Moreover, the expected larger cell sizes of polyploids vs. diploids might allow for more and larger mitochondria, which in turn give rise to opportunities for compensatory regulation at increasing levels of organization including the number of organelle genome copies per organelle, the number of transcripts/proteins produced from each organelle genome copy, and even the number of OXPHOS complexes formed per unit area of the mitochondrial inner membrane^97^. Across evolutionary time, if mitochondrial performance is indeed a critical component of fitness in *P. antipodarum*, as it is for other animal taxa^1–5^, asexual lineages with decreased mitochondrial performance might be evolutionarily short lived and difficult to sample. Combined with the strong lake effect observed here, it is clear that only extensive within-lake sampling combined with integrative approaches that span levels of biological organization will be capable of elucidating mechanisms of mitochondrial compensation in asexual lineages of *P. antipodarum*.

The nature of our sampling design means that the lake effects observed here could be the result of either genetic or environmental forces. In particular, these effects are largely consistent with our current understanding of *P. antipodarum* population genetic structure^56,59,61^, in which lakes and other geographic features of New Zealand (e.g., the Southern Alps^56^) act as major barriers to gene flow among *P. antipodarum* subpopulations. Previous work investigating mitochondrial function in lab-cultured iso-female asexual lineages of *P. antipodarum* (i.e., snails that are asexual descendants of a single mother collected from the wild and isolated in the lab) has identified heritable variation for mitochondrial, metabolic, and behavioral phenotypes^27^. However, because the snails used in this study were born and lived to reproductive maturity in the wild and subsequently acclimated to laboratory conditions, it is possible that the lake effects observed here might also be the result of phenotypic plasticity for metabolic rate. One tractable angle for investigating the contributions of genetic vs. environmental variation in mito-nuclear genotype to variation in mitochondrial performance can take advantage of extensive mito-nuclear discordance observed in asexual lineages of *P. antipodarum*^59^. Because asexual lineages stably transmit mito-nuclear genotypes across generations, asexual lineages that have trapped reciprocal combinations of mitochondrial haplotypes with different nuclear genotypes can be used to test whether mito-nuclear epistatic variation contributes to mitochondrial function and ultimately, organismal fitness.

Maternal transmission of mitochondrial genomes precludes inheritance of all mitochondrial mutations originating in males, with two primary consequences for the efficacy of selection on the mitochondrial genome in sexual taxa. First, genes in the mitochondrial genome of sexually reproducing lineages experience a reduction in the effective population size relative to nuclear genes due to both the absence of recombination and uniparental inheritance. Second, mutations in the mitochondrial genome that have sex-specific effects only experience effective natural selection in females^98,99^. This latter phenomenon is predicted to result in the accumulation of mutations that are neutral or beneficial in females but deleterious in males^100^. The lack of widespread evidence for mitochondrial mutations with sex-specific fitness effects (but see^35,101,102^) may point to mechanisms that prevent the spread of male-specific deleterious mutations in mitochondrial genomes (*e.g.,* paternal leakage^103^, inbreeding^104^, kin selection^104^, and/or nuclear-encoded restorers of male function^42,105^). Asexually produced males, as are occasionally produced in *P. antipodarum*^60^, represent worst-case scenarios for male-specific mitochondrial performance because their nuclear and mitochondrial genomes have been “trapped” in females for generations but are now being expressed in a male context. Because selection against male-harming mutations is therefore completely ineffective in asexual lineages, asexual males are expected to exhibit particularly poor mitochondrial performance. We found that male *P. antipodarum* have higher mitochondrial membrane potential than female counterparts, leading us to speculate that male *P. antipodarum* do not suffer from sex-specific mutations that decrease their ability to generate a proton motive force. Mitochondrial membrane potential depends upon protons pumped by the OXPHOS complexes that are comprised of both nuclear and mitochondrial gene products (*i.e.,* complexes I, III, and IV^32,33^). The implications are that we should expect to detect reduced mitochondrial membrane potentials if male-harming mutations have accumulated in *P. antipodarum* mitochondrial genomes. One of the most intriguing findings presented here is that asexual males appear to be driving this pattern of elevated mitochondrial membrane potential to a greater extent than sexual males. However, because we do not know the relationship between mitochondrial membrane potential and fitness, we cannot yet determine whether this sexual dimorphism in mitochondrial membrane potential supports or refutes the presence of male-harming mitochondrial mutations in *P. antipodarum*. Still, the marked increase in mitochondrial membrane potential among asexually produce males offers a promising avenue for investigating the prevalence of evolutionary dynamics of male-harming mutations.

## CONCLUSIONS

We found strong effects of temperature on oxygen consumption, righting behavior, and emergence time, lake effects on three out of four mitochondria-related phenotypes, a difference among sexual and asexual individuals in one behavioral assay, and a sex-specific difference in mitochondrial membrane potential. These results indicate that local conditions may play a major role in modulating mitochondrial function in *P. antipodarum*. The strong lake effects observed demand more extensive intra-lake sampling, in order to better test for the effects of reproductive mode and sex on mitochondrial and organismal performance in *P. antipodarum*. Nevertheless, with the current sampling, we were not able to detect a global effect of asexual reproduction to decrease performance. This last finding points to physiological or genetic compensation for the putative accumulation of deleterious mutations in mitochondrial genomes in asexual *P. antipodarum*. Together, our results establish *P. antipodarum* as a model system for evaluating the consequences of sexual reproduction for mitochondrial function and evolution and for evaluating the strength and efficacy of selection against male-harming mitochondrial mutations in sexual populations.

## DATA ACCESSIBILITY

Oxygen consumption, behavioral, and mitochondrial membrane potential data have been posted to https://github.com/jsharbrough/Potamo_mt_function and will be deposited in Dryad (https://doi.org/10.5061/dryad.z08kprr8w) and FigShare (https://doi.org/10.6084/m9.figshare.11782128).

## ACKNOWLEDGEMENTS

This work is dedicated in loving memory to J.J. Neiman-Brown, whose smile and curiosity give inspiration to us all. This manuscript was written in part during the Biology Department Weekly Writing Retreat at Colorado State University. Flow cytometry was performed at UI’s Flow Cytometry Facility. We thank two anonymous reviewers for comments on an earlier version of this manuscript, and Laura Bankers, Kaitlin Hatcher, Katelyn Larkin, and Kyle McElroy for snail collections. We thank the organizers, especially Karen Burnett, and all the participants of the SICB 2020 Building Bridges from Genome to Phenome: Methods, Molecules, and Models symposium for their participation in the symposium and helpful feedback on this work.

---

**Table S1.**
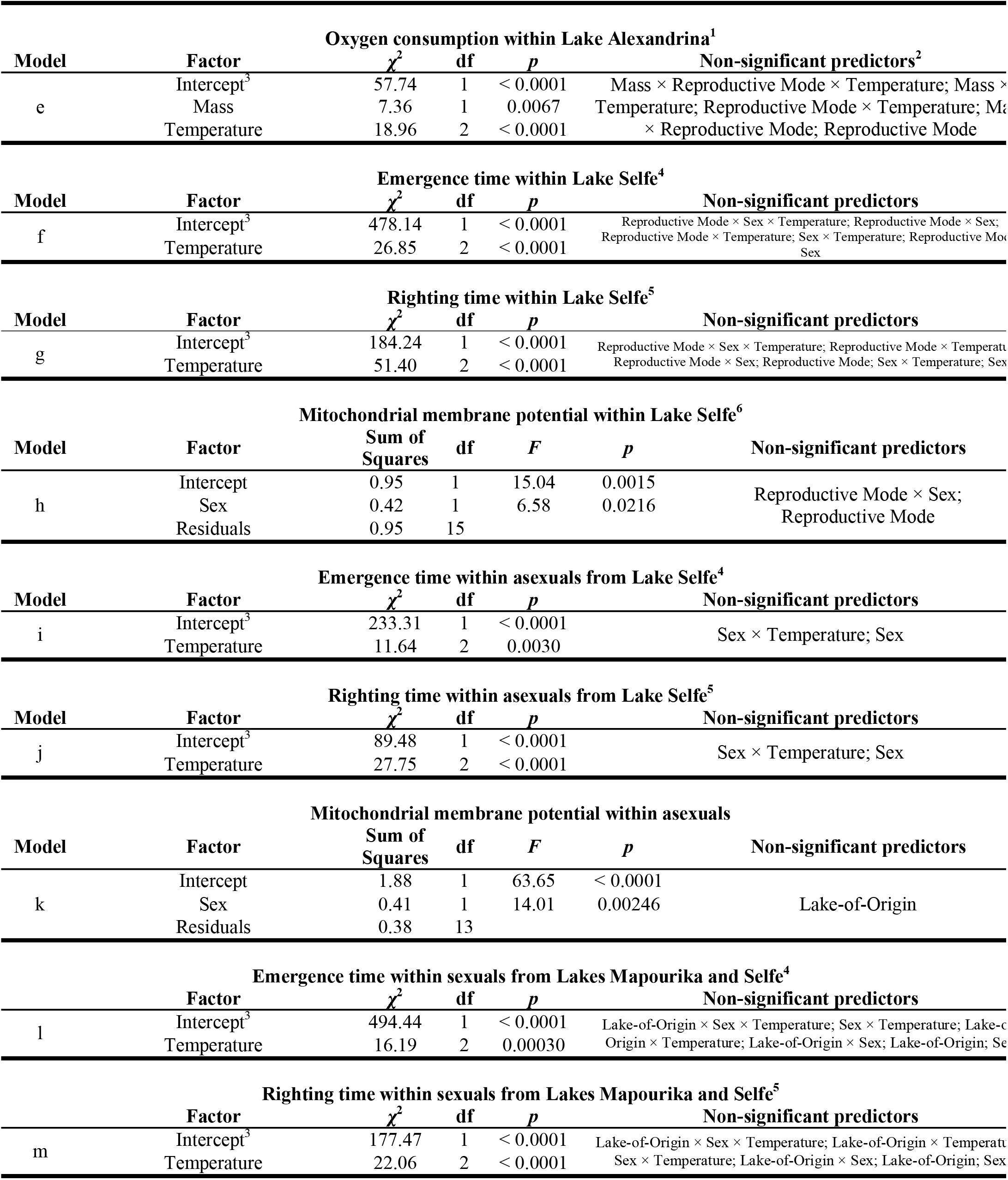
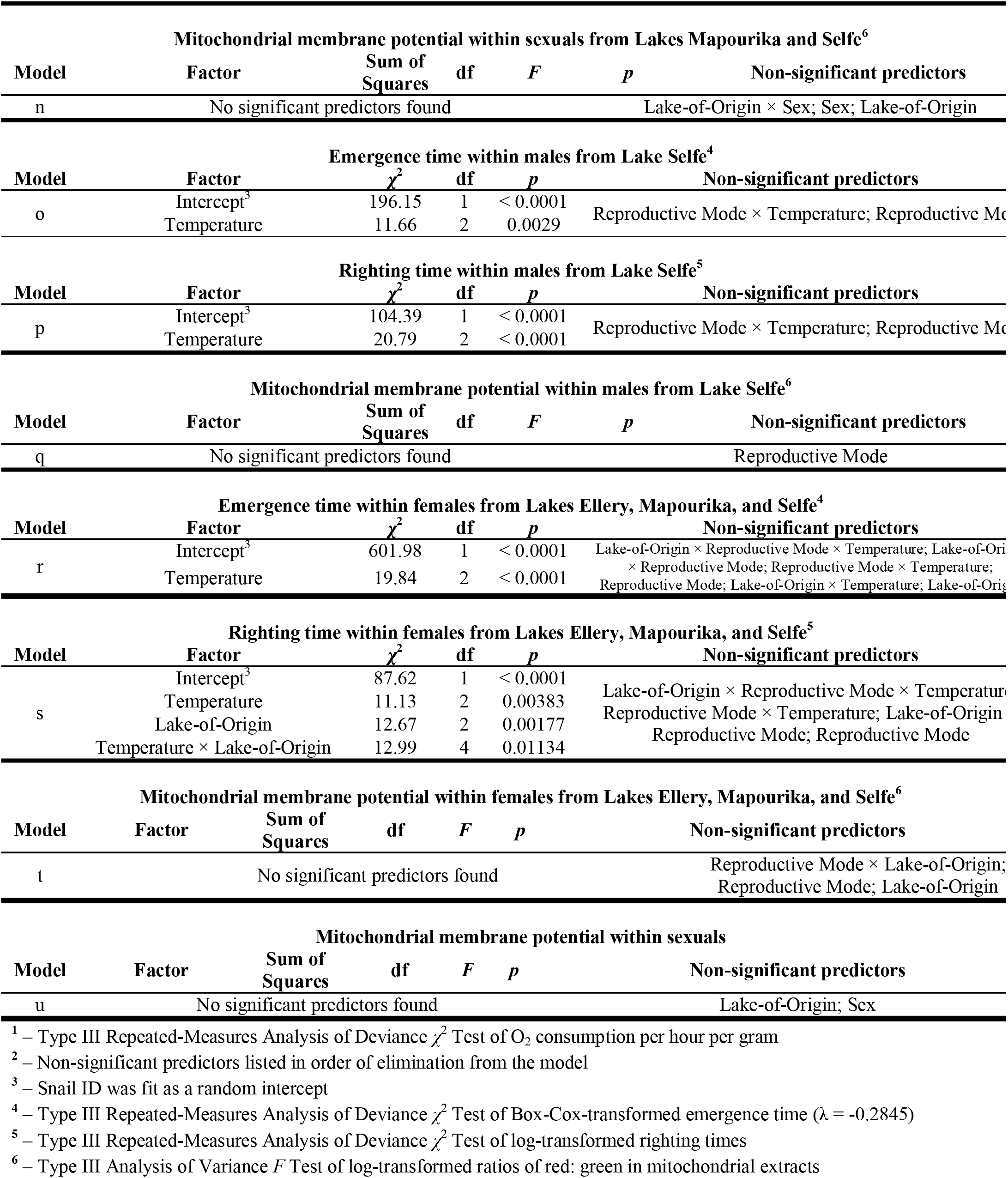
Fixed-effect only and mixed-effects models of select predictors on oxygen consumption, righting time, emergenc time, and mitochondrial membrane potential nested within lakes.

**Figure S1.**
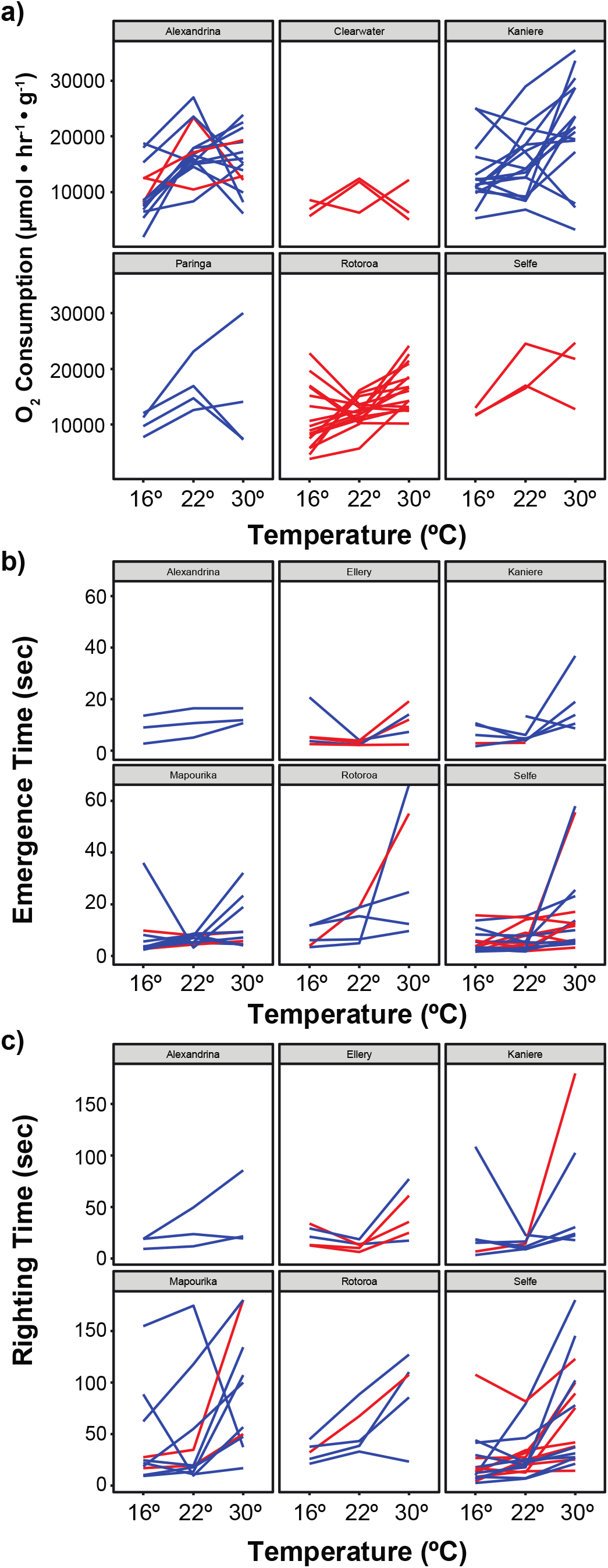
Individual snail reaction norms across lakes in response to temperature stress. Lines in each box (lake) represent individual reaction norms collected across three temperatures (16°C, 22°C, and 30°C) for each of three mitochondria-related phenotypes. Red lines represent asexual individuals and blue lines represent sexual individuals. **a)** Oxygen consumption per hour per gram. **b)** Time to emergence following a startling stimulus. **c)** Righting time after being flipped.

**Figure S2.**
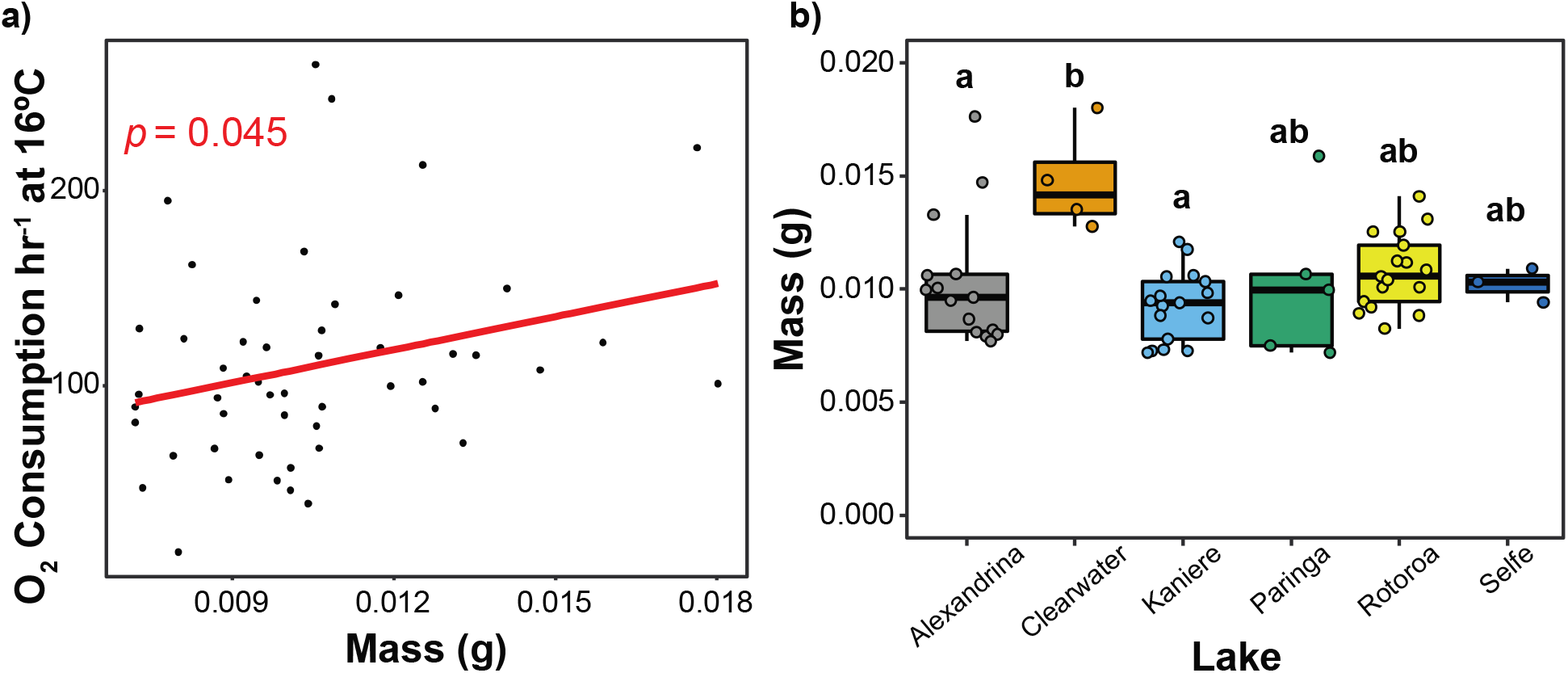
Snail mass and oxygen consumption. **a)** Snail mass is positively correlated with O_2_ consumption at 16°C. **b)** Boxplot showing all datapoints (filled circles) depicting snail mass across lakes. Snail mass varies across snails sampled from different lakes. Lowercase letters above boxplots indicate statistical groupings at the α=0.05 level based on pairwise t-tests, correcting for multiple comparisons using the Holm procedure.

**Figure S3.**
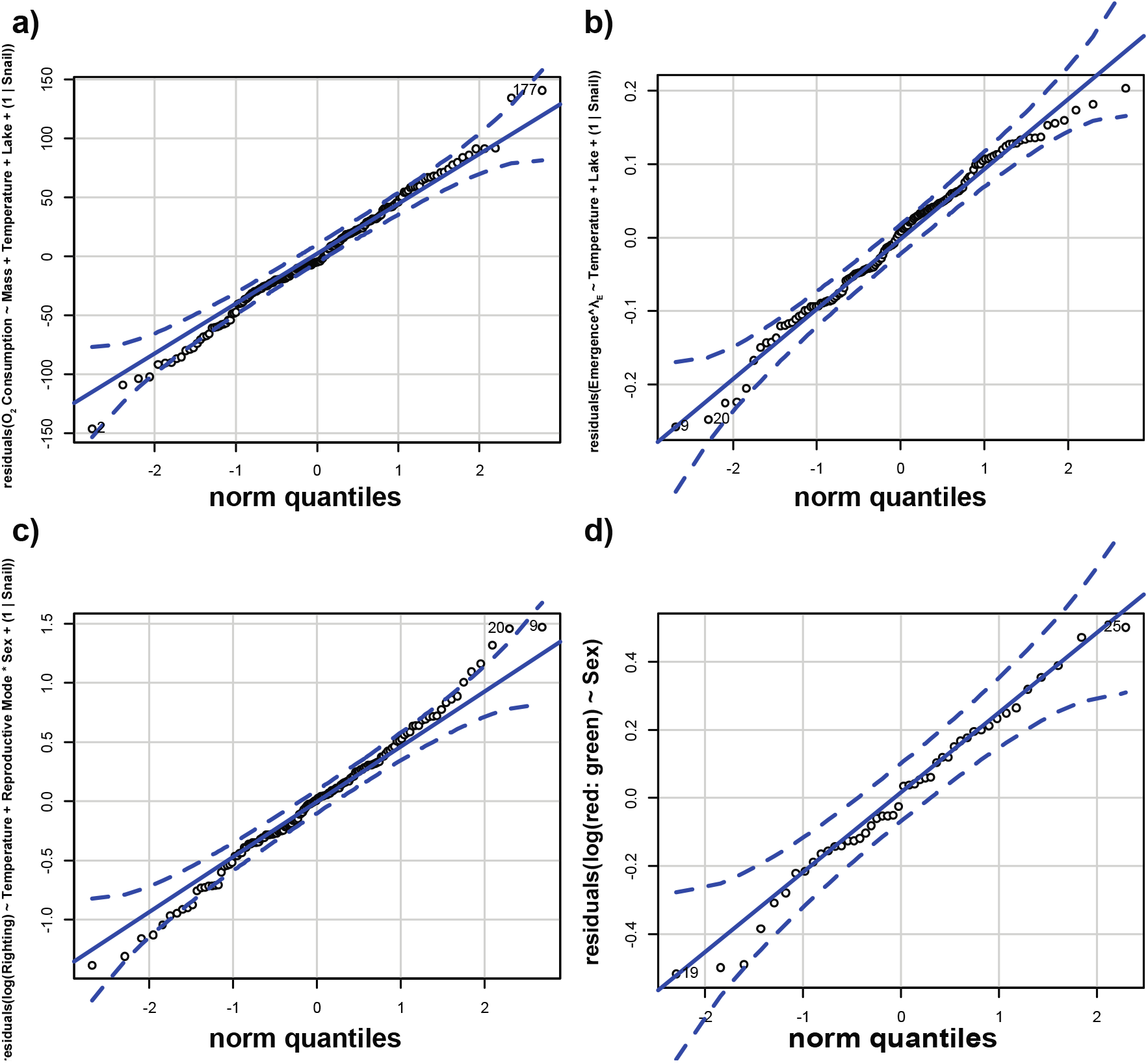
QQ plots used to identify appropriate statistical transformations. Norm quantiles plotted against: **a)** Oxygen consumption model residuals. **b)** Power-transformed emergence time model residuals (λ_E_ = −0.2845). **c)** Log-transformed righting time model residuals. **d)** Log-transformed ratio of red: green fluorescence following treatment with JC1.

